# Pinch2 is a novel regulator of myelination in the Central Nervous System

**DOI:** 10.1101/2022.03.04.483000

**Authors:** J Paes de Faria, RS Vale-Silva, R Fässler, HB Werner, JB Relvas

## Abstract

The extensive morphological changes of oligodendrocytes during axon ensheathment and myelination involve assembly of the Ilk-Parvin-Pinch (IPP) heterotrimeric complex of proteins to relay essential mechanical and biochemical signals between integrins and the actin cytoskeleton. Binding of Pinch 1 and 2 isoforms to Ilk is mutually exclusive and allows the formation of distinct IPP complexes with specific signaling properties. Using tissue-specific conditional gene ablation in mice, we reveal an essential role for Pinch2 during central nervous system myelination. Unlike *Pinch1*-gene ablation, loss of *Pinch2* in oligodendrocytes results in hypermyelination and in the formation of pathological myelin outfoldings in white matter regions. These structural changes concurred with inhibition of Rho GTPases RhoA and Cdc42 activities and phenocopied aspects of myelin pathology observed in corresponding mouse mutants. We propose a dual role for Pinch2 in preventing excess of myelin wraps through RhoA-dependent control of membrane growth and in fostering myelin stability via Cdc42-dependent organization of cytoskeletal septins. Together, these findings indicate that IPP-containing Pinch2 is a novel critical cell-autonomous molecular hub ensuring synchronous control of key signaling networks during developmental myelination.

**Summary statement:** Pinch proteins are core components of a ternary protein complex comprising Ilk and Parvin (IPP). This work identifies Pinch2 as key regulator of the formation and maturation of CNS myelin.

## Introduction

In the central nervous system (CNS), oligodendrocytes (OL) form myelin, a specialized cytoplasmic membrane growing around axons that ensures fast and effective propagation of action potentials along axons (Nave and Werner, 2014, Snaidero and Simons, 2017). The molecular signals regulating myelination derive, at least in part, from instructive cues originated by the extracellular matrix (ECM) and transduced by integrin transmembrane receptors. This signaling is initiated by ligand occupation and clustering of integrin subunits in topographically discrete regions of the membrane, and mediated through molecular scaffolds, such as the heteroternary IPP complex in focal adhesions (Legate et al., 2006, Wickstrom et al., 2010, Qin and Wu, 2012, Wu, 2004). In association with other molecules – such as paxillin, talin, vinculin, and tensin - this protein complex links the ECM with the actin cytoskeleton by regulating the activity of downstream molecules, including Rho GTPases, Profilin-1, Jab-1 or N-Wasp (Feltri et al., 2016). The IPP complex contains Integrin Linked Kinase, Ilk, the adaptor protein five LIM (Lin-1, Isl-1, Mec-3) domain-containing protein Pinch (Particularly Interesting Cys-His-rich protein), and the F-actin binding proteins Parvins. Pinch and Ilk interaction, which occurs prior to the formation of cell-ECM adhesion complexes in the cell membrane independently of adhesion signals (Zhang et al., 2002b), is necessary to prevent proteasome-mediated degradation of the IPP complex (Fukuda et al., 2003). In mammalian cells, IPP function depends on the mutually exclusive binding (Li et al., 1999, Zhang et al., 2002a) of Pinch1 (Rearden, 1994, Tu et al., 1999) or Pinch2 (Hobert et al., 1999, Zhang et al., 2002a), and forms a signaling hub controlling critical cell functions as cell – ECM contact and adhesion, organization of the cytoskeleton, survival, proliferation and migration in various cellular contexts (Legate et al., 2006). Encoded by distinct genes-*Lims1* and *2* - the two proteins have high sequence similarity (Braun et al., 2003), and are co-expressed in various tissues (Rearden, 1994, Braun et al., 2003). Pinch1 is highly and mostly exclusively expressed in early embryonic stages, whereas Pinch2 is detected from the midgestational stage (Braun et al., 2003). The two Pinch isoforms have different binding partners (Kadrmas et al., 2004, Dougherty et al., 2005, Vaynberg et al., 2005, Velyvis et al., 2003, Wiesner et al., 2005, Guo et al., 2019) whose recruitment likely confers signaling specificity to the IPP complex. In the peripheral nervous system, Ilk is required for the developmental radial sorting of axons out of bundles through negative regulation of Rho/Rho kinase signaling (Pereira et al., 2009). In oligodendrocytes, Ilk is required for laminin-2-induced formation of myelin-like membrane formation (Chun et al., 2003) and regulates process extension by regulating OL actin cytoskeletal dynamics (O’Meara et al., 2013, Michalski et al., 2016). *In vivo*, Ilk ablation disturbs OPC proliferation and differentiation (Hussain and Macklin, 2017). However, as it prevents assembly of the IPP complex it also leads to the degradation of the other IPP components making it impossible to dissect the exact role played by Pinch1 and 2 in OL development. Here, we ablated either *Lims1* /*Pinch1* or *Lims2* /*Pinch2* in the mouse OL lineage to investigate their specific functions during myelination. Our results support an essential function of Pinch2 protein in controlling the development and structure of myelin sheath.

## Results

### Pinch1 and Pinch2 expression is developmentally regulated in oligodendrocytes

We examined the expression of the two Pinch proteins in the mouse optic nerve (ON), an OL-enriched tissue that follows a well-established spatiotemporal sequence of events during development (Dangata and Kaufman, 1997, Foran and Peterson, 1992) and in which axons are uniformly aligned. In mice, OL precursor cells (OPC) migrate from the floor of the third ventricle throughout the developing nerve until around the day of birth; at postnatal day 5 (P5) most OPCs are differentiating and myelination of axons initiates around P7. While Pinch1 expression is already detected at P5 **(Fig. 1A,B)**, Pinch2 expression is detected later at P15 **(Fig. 1A,C)**. These protein profiles reflect the abundance of their respective RNA molecules in the developing myelinating glia (Zhang et al., 2014). To understand Pinch1 and Pinch2 function in oligodendrocytes, we generated conditional *Pinch1* and *Pinch2* single cKO by *in vivo Cre*-mediated recombination. Mouse lines carrying *Lims1* /*Pinch1* or *Lims2* /*Pinch2* with relevant *floxed* allelic regions (Li et al., 2005, Stanchi et al., 2005) (**supplementary material Fig. S1A)** were crossed with a *Cnp* driven *Cre* mouse line (*Cnp*^*Cre/+*^), where *Cre* is expressed in differentiating myelinating glia from embryonic day 13 (Lappe-Siefke et al., 2003). The resulting conditional knockout (cKO) progenies *Cnp*^*Cre/+*^ *Pinch1*^*flox/flox*^ (hereafter referred to as *Pinch1* cKO or *P1* cKO) or *Cnp*^*Cre/+*^ *Pinch2*^*flox/flox*^ (hereafter referred to as *Pinch2* cKO or *P2* cKO) and the respective littermates, *Pinch1*^*flox/flox*^ and *Pinch2*^*flox/flox*^ (controls, CTR) were born at the expected Mendelian frequency and displayed no obvious developmental or motor deficits. Conditional recombination of either *floxed Pinch1* or *Pinch2* alleles resulted in significantly decreased protein levels in lysates obtained from P15-days old ONs **(Fig. 1D)**. The residual levels of Pinch detected in the blot are likely due to the presence of astrocytes and low numbers of unrecombined oligodendrocytes in the tissue (Zhang et al., 2014).

**Figure 1.**
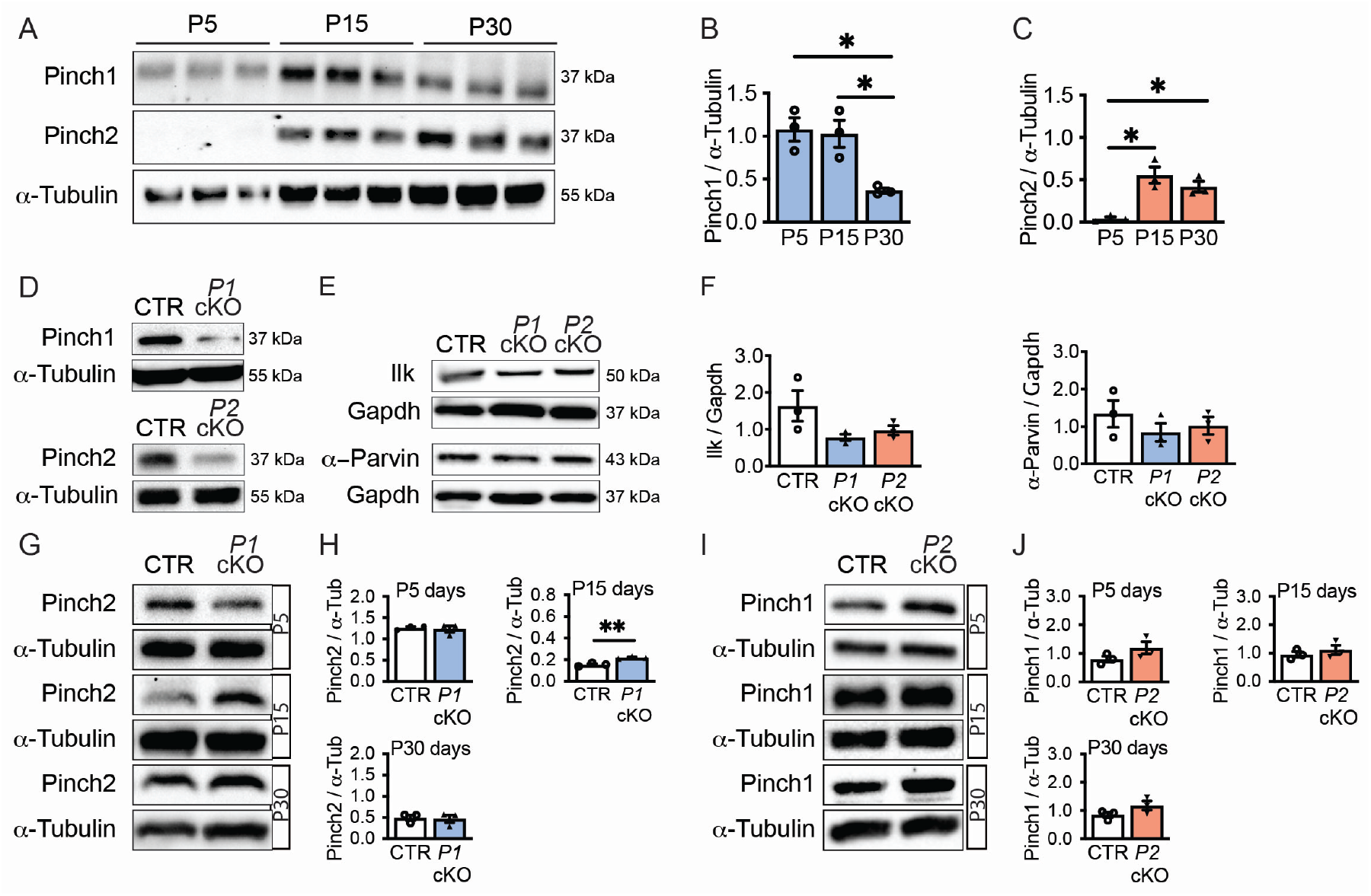
Pinch proteins are expressed differently in the developing optic nerve. **(A)** Western blot analysis of optic nerve (ON) homogenates derived from P5, P15 and P30 wild-type mice (C57BL/6). **(B, C)** Densitometric analysis of Pinch1 and Pinch2 immunoreactive signals upon normalization with a-Tubulin. Statistical differences detected between P5-P30 (*P* =0.019) and P15-P30 (*P* =0.019) for Pinch1 and P5-P15 (*P* =0.005) and P5-P30 (*P* =0.024) for Pinch2. Analysis used one-way ANOVA: F(2,6)=10.52, *P* =0.01 and F(2, 6)=15.21 *P* =0.004, followed by a *Holm-Sidak post hoc* test, *n*=3. **(D)** Western blot analysis of Pinch1 and Pinch2 in *Pinch1* cKO (*Cnp*^*Cre/+*^:*Pinch1*^*flox/flox*^, *P1* cKO,) and *Pinch2* cKO (*Cnp*^*Cre/+*^:*Pinch2*^*flox/flox*^, *P2* cKO) mice, respectively and the correspondent controls (*Pinch1*^*flox/flox*^ and *Pinch2*^*flox/flox*^, CTR) at P15. **(E, F)** No significant changes detected in the levels of Ilk and a-Parvin in *Pinch* cKOs when compared with CTR. One-way ANOVA F(2, 6)=3.11 (*P* =0.11) in Ilk and F(2, 6)=0.76 (*P* =0.5) in a-Parvin (*n*=3). **(G-J)** Immunoblot analysis of ONs from *Pinch* cKO and CTR mice at P5, P15 and P30 show unchanged levels of the untargeted *Pinch*, except a transient upregulation of Pinch2 in *P1* cKO (ON, P15). Unpaired two-tailed Student’s *t*-test, *n*=3 **(H, J)**. Bars represent means±s.e.m. (**B, C, F, H, J)**.

The IPP protein complex arises from the association of different proteins, including the exclusive binding of either Pinch1 or Pinch2 to Ilk (Li et al., 1999, Zhang et al., 2002a). Thus, the loss of one Pinch protein does not preclude the assembly of an alternative complex with the existing Pinch, Ilk and Parvin (Fukuda et al., 2003). By immunoblot we confirmed that in P15 ON the expression levels of Ilk and a-Parvin in single individual *Pinch* cKO were not significantly different from controls **(Fig. 1E,F)**. This establishes that in the presence of either Pinch, the IPP complex of proteins is formed. Targeting one *Pinch* and depleting its expression did not result in a compensatory increase in the expression of the untargeted *Pinch* counterpart **(Fig. 1G–J)**, with the conspicuous exception of P15 *Pinch1* cKO ONs, in which Pinch2 expression was transiently increased (0.15±0.009 *versus* (*vs*) 0.22±0.007, *P* =0.005; **Fig. 1G,H)**, which may be due transient accumulation of unbound Ilk molecules and increase in Pinch2 stability (Stanchi et al., 2005). Altogether these results show that in oligodendrocytes, the IPP complex is formed by either Pinch1 or 2 proteins at distinct developmental stages possibly to exert different context-specific functions.

### Axon contact and ensheathment are not affected by the loss of either Pinch1 or Pinch2

To gain insight into the individual contribution of Pinch1 and 2 in developmental myelination, we first examined the myelin ultrastructure of white matter tracts in the optic nerve, brain (midsagittal sections of the corpus callosum) and spinal cord at P15, a relatively early stage of developmental myelination, by transmission electron microscopy (TEM). At P15, despite the loss of either *Pinch1* or *Pinch2*, oligodendrocytes were able to efficiently myelinate axons in these different anatomical regions **(Fig. 2A)**. In line with this, no significant differences were found in the numbers of contacted axons (*P* =0.77, *n*=5 for *P1* cKO; *P* =0.42, *n*=3/4 for *P2* cKO; **Fig. 2B**) or in the relationships between axon diameter and myelin sheath thickness (*g*-ratio) (0.842±0.001 *vs* 0.846±0.003, *P* =0.21, *n*=3 in *P1* cKO; 0.849±0.005 *vs* 0.854±0.005, *P* =0.27, *n*=3 in *P2* cKO; **Fig. 2C**) in the ON of *Pinch* cKOs and their control littermates. These observations suggest that the formation of an IPP complex by either Pinch is sufficient to support the onset and the early stages of CNS myelination. Ilk (Pereira et al., 2009) and Pinch1 (Eke et al., 2010, Fukuda et al., 2003) were previously described to be required for the activation of Akt, which is also an essential modulator of OPC survival *in vitro* (Flores et al., 2000). However, we did not detect significant changes in the levels of Akt phosphorylation in lysates obtained from ON of P15 *Pinch1* cKO compared to those from controls (*P* =0.26, *n*=3; **supplementary material Fig. S2A**). Similarly, the activity of p38 mitogen-activated protein kinase (MAPK), which is also regulated via IPP (Smeeton et al., 2010), remained unchanged in ON lysates from either *Pinch* (*P* =0.60, *n*=3 in *P1* cKO; *P* =0.68, *n*=4 in *P2* cKO; **supplementary material Fig. S2B**). In contrast, the activity of the MAP kinases Erk1/2 and Jnk were specifically and transiently upregulated in P15 *Pinch1* cKO (p-Erk: 0.82±0.08 *vs* 1.14±0.03, *P* =0.018, *n*=3; p-Jnk: 0.24±0.02 *vs* 0.42±0.05, *P* =0.039, *n*=3; **supplementary material Fig. S2C,D**). Because sustained activation of Erk1/2 increases the numbers of oligodendrocyte precursors (OPCs) (Ishii et al., 2013), we next examined OPC/OL cells in P15 ONs of both *Pinch1* and *Pinch2* cKOs using the *bone fide* lineage marker, Olig2. We observed a transient increase in Olig2-positive cells in the ON of *Pinch1* cKO (49.8±2.4 *vs* 59.8±2.2, *P* =0.036, *n*=3; **supplementary material Fig. S2E**) but not in *Pinch2* cKO (*P* =0.64, *n*=3; **supplementary material Fig. S2F**), which is consistent with the increase in Erk1/2 activity in *Pinch1* cKO. OL-lineage expansion coexisted with stable numbers of CC1-positive mature oligodendroglia in *Pinch1* cKO (34.2±2.4 *vs* 40.3±0.7, *P* =0.07, *n*=3; **supplementary material Fig. S2E**), suggesting that OPC/OL population growth is paralleled by steady OL differentiation and increased cell death. This was confirmed by the presence of significantly higher numbers of Olig2-positive cells which were also TUNEL-positive in the ON of *Pinch1* cKO (4.9±0.6 *vs* 8.9±0.5, *P* =0.0089, *n*=3; **supplementary material Fig. S3A**), compared with controls at P30. As Pinch1 inhibits Jnk-mediated apoptosis in mouse primitive endoderm cells (Montanez et al., 2012), and its activation is increased in lysates obtained from *Pinch1* cKO ONs, it is conceivable that a Pinch1-dependent mechanism regulates OPC/OL pool via Jnk-mediated cell apoptosis. Concomitant with the transient variation in the numbers of Olig2 positive cells, both Erk1/2 and Jnk activities in *Pinch1* cKO return to control levels by P30 (P-Erk: 1.2±0.1 *vs* 1.1±0.2, *P* =0.8, *n*=4; P-Jnk: 1.6±0.1 *vs* 1.4±0.1, *P* =0.3, *n*=3; **supplementary material Fig. S3B,C**).

**Figure 2.**
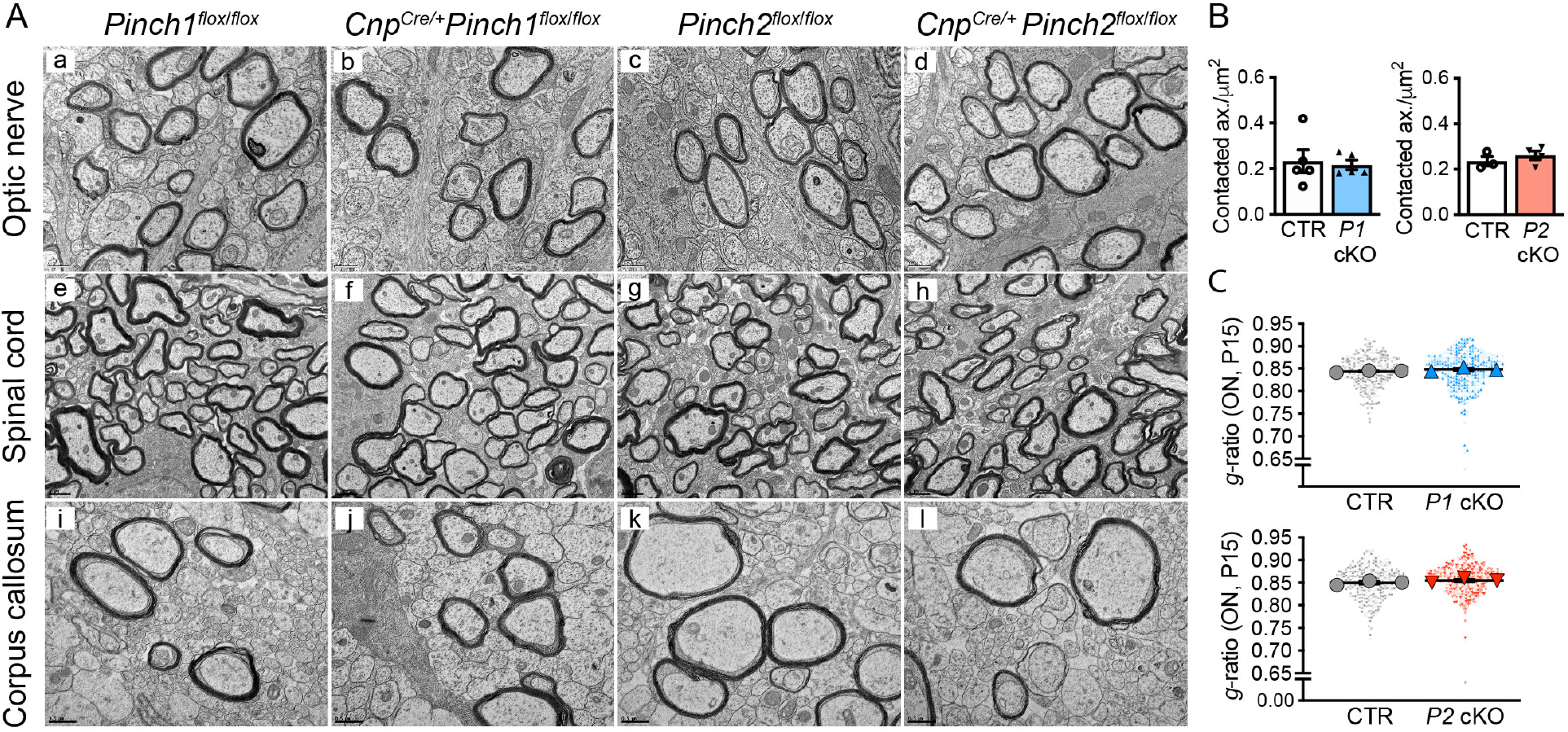
*Pinch* depletion does not prevent axon contact and ensheathment. **(A)** Representative electron micrographs of cross-sections of the optic nerve (ON, a-d), spinal cord (e-h) and corpus callosum (CC, i-l) from conditional *Pinch* KO mice, *Pinch1* (*Cnp*^*Cre/+*^ *Pinch1*^*flox/flox*^; *P1* cKO in b, f, j) and *Pinch2* (*Cnp*^*Cre/+*^ *Pinch2*^*flox/flox*^; *P2* cKO in d, h, l) and from the respective controls (CTR in a, e, i, and in c, g, k) at P15. Scale 0.5 μm (ON and CC), and 1 μm (SC). **(B)** Similar number of contacted and/or myelinated axons in the ON from *Pinch* cKOs and CTR mice at P15. Bars represent means±s.e.m.. **(C)** No differences in *g*-ratio in cKO compared with the respective CTR (measured in ONs at P15). Beeswarm superplots show mean *per* animal used to calculate average, SEM, and *P* values. Statistical analysis used unpaired two-tailed Student’s *t*-test in **(B)** and **(C)**, *n*=3/4 (in **(C)**, each *n* represents at least 300 myelinated axons analyzed *per* genotype).

These findings suggest that in the early developing ON, Pinch1 - but not Pinch2 - is involved in the control of OPC/OL population homeostasis, through MAPK-dependent signaling. Because independent ablation of Pinch1 and 2 had not an obvious impact in axon contact or ensheathment, our results suggest that Pinch proteins are competent in establishing IPP during the onset and early stages of myelination.

### Loss of Pinch2 results in myelin overgrowth and formation of redundant myelin

We next examined the myelin ultrastructure in cross-sections from ON, spinal cord, and corpus callosum in young adult mice (P30), a period of active myelination. We observed that in these distinct anatomical regions, the myelin sheaths of *Pinch2* cKO axons were thicker compared to that of control or *Pinch1* cKO axons **(Fig. 3A)**. This was reflected by the significantly lower *g*-ratios found in *Pinch2* cKO (0.811±0.012 *vs* 0.773±0.0051, *P* =0.015, *n*=3/5; **Fig. 3B**, **supplementary material Fig. S4A**), while in *Pinch1* cKO *g*-ratios remained close to control values (0.802±0.008 *vs* 0.806±0.003, *P* =0.62, *n*=3). In both cKOs, the number of myelinated axons was similar to controls (*P* =0.9, *n*=3 in *P1* cKO and *P* =0.4; *n*=3/5 in *P2* cKO; **supplementary material Fig. S4B**). Morphometric analysis of myelinated axons from the ventral white matter of the spinal cord of the *Pinch* cKOs, showed that only the depletion of *Pinch2* resulted in thicker myelin sheaths (0.791±0.002 *vs* 0.750±0.013, *P* =0.035, *n*=3; **supplementary material Fig. S4C**). We confirmed myelin thickening excluding the contribution of any variation in periaxonal compartment (**Fig. 3C**, *P* =0.97, *n*=3/5; **supplementary material Fig. S4D,E**; 0.854±0.002 *vs* 0.832±0.004, *P* =0.036; *n*=3-5) (Goebbels et al., 2010) or changes in the periodicity of compact myelin lamellae (*P* =0.53, *n*=3; **supplementary material Fig. S4F**), which supports the view that thicker myelination resulted from extra myelin wraps. Although the formation of aberrantly thick myelin sheaths can result from sustained activation of the Erk1/2 MAPK pathway (Ishii et al., 2013, Jeffries et al., 2016), this is likely not the case, as we did not detect significant changes in the levels of phospho Erk1/2 between *Pinch2* cKO and control nerves (*P* =0.76, *n*=3; **supplementary material Fig. S4G**). In addition, myelin overgrowth in *Pinch2* cKO is unlikely the result of hyperactivation of Akt (Flores et al., 2008) or the negative regulation by PTEN (Goebbels et al., 2010) (Akt: *P* =0.36, *n*=3; PTEN: *P* =0.29, *n*=3; **supplementary material Fig. S4H,I**). The increase in the number of myelin sheaths-overmyelination-may result in focal hypermyelination or emergence of myelin abnormalities around myelinated axons. In addition to myelin overgrowth, EM analysis of optic nerve, spinal cord, or corpus callosum at P30 also revealed the presence of redundant membrane outwards loops (myelin outfoldings, indicated by red arrowhead in **Fig. 3A**, **Fig. 3D, supplementary material Fig. S5A**) and isolated “myelin-like” accumulations (myelin “whorls”, indicated by blue arrows in **Fig. 3D** and **supplementary material Fig. S5A)** in *Pinch2* cKO. Although noticeable at P30 **(supplementary material Fig. S5A–C)**, morphometric analysis revealed that these myelin defects were significantly more abundant in P90 *Pinch2* cKO nerves compared to their age-matched controls (*P* =0.0007, *n*=6 in ON; *P* =0.024, *n*=3-4 in SC; **Fig. 3 D-F**). To eliminate the possibility that the loss of one *Cnp* allele *per se* was affecting the myelin ultrastructure, we quantified the frequency of either myelin outfoldings and whorls in ultrathin sections of the spinal cord (SC) from *Pinch2*^*flox/wt*^ *Cnp*^*Cre/+*^ mice and concluded that their frequency was similar to that of *Pinch2*^*flox/flox*^ control mice (*P* =0.4, *n=3*; **supplementary material Fig. S5D-E**). Together, these results strongly suggest that Pinch2-mediated signaling is required for normal myelination of CNS tracks. Aberrant myelin profiles in the CNS are often associated to anomalous axoglial adhesion (Djannatian et al., 2019, Elazar et al., 2019). These contacts are essential to form paranodal axoglial junctions at the end of each myelin segment, where a cytoplasm-filled structure (or paranode loop) flanks the node of Ranvier. We examined paranode loops in longitudinal sections of the ON and ventral SC of P90 *Pinch2* cKO and control mice. While in controls most of these structures were geometric clustered at paranodes in an orderly fashion and contacting the axolemma **(supplementary material Fig. S5F)**, white arrowheads), in *Pinch2* cKOs these structures were often disorganized and did not establish appropriate contacts with the axon surface (white arrowheads). Defective contacts were better visualized at high magnification and we observed that in some paranodes, the loops were piled up in disarray order and often did not contact the axon but sit on top of compacted myelin. This was particularly evident in the outer loops **(supplementary material Fig. S5F**, asterisks). We next measured the length of nodes of Ranvier in the spinal cord and optic nerve of *Pinch2* cKO and control adult mice **(supplementary material Fig. S5G**, black dashed line). Strikingly, we found that in both anatomical regions, node length was increased (0.79±0.03 *vs* 1.06±0.06, *P* =0.018, *n*=3 in SC; 0.83±0.03 *vs* 1.04±0.06, *P* =0.033, *n*=3 in ON) and distributed over a broader range in *Pinch2* cKO compared to controls **(supplementary material Fig. S5H,I)**. These alterations were not significantly correlated with either the node diameter or axon diameter at the paranode **(supplementary material Fig. S5J–M)**. These results suggest that OL-specific loss of *Pinch2* disrupts the node architecture potentially through the disintegration of critical axo-glia contacts, supporting a Pinch2-dependent function in myelin assembly and stability.

**Figure 3.**
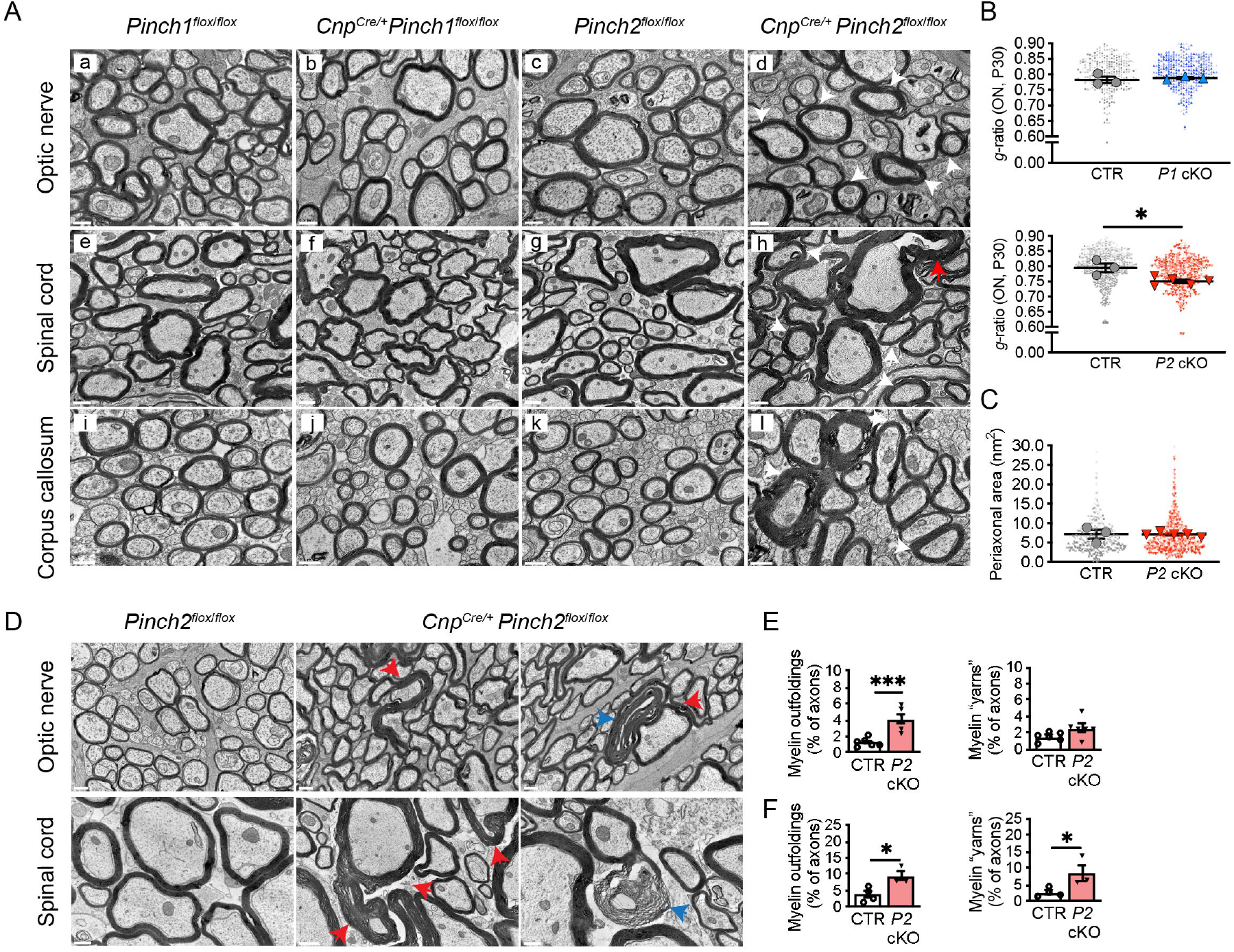
OL-specific loss of *Pinch2* leads to myelin membrane overgrowth and formation of myelin outfoldings. **(A)** Representative electron micrographs of cross-sections of the optic nerve (ON, a-d), spinal cord (SC, e-h) and corpus callosum (CC, i-l) from conditional *Pinch1* KO (*Cnp*^*Cre/+*^ *Pinch1*^*flox/flox*^, *P1* cKO in b, f, j) and conditional *Pinch2* KO (*Cnp*^*Cre/+*^ *Pinch2*^*flox/flox*^, *P2* cKO in d, h, l) and the respective controls (CTR in a, e, i, and c, g, k) at P30. Examples of axons with thicker myelin (white arrows) and myelin aberrations (red arrows) are indicated. Scale bars: 0.5 μm, except SC (1 μm). **(B)** Lower *g*-ratio in *P2* cKO compared with the respective CTR. In contrast, no significant differences found between *P1* cKO and CTR. Beeswarm superplots show the means *per* animal used to calculate average, SEM, and *P* values (*n*=3/5). At least 300 axons *per* animal were measured (smaller symbols). **(C)** Periaxonal area is similar in *P2* cKO and CTR mice (ON). Mean value *per* animal (used to calculated average and SEM) is represented as a larger symbol in the beeswarm superplot (*n*=3/5). **(D)** Representative electron micrographs of cross-sections of ON and SC from *P2* cKO and CTR littermates at P90. Myelin outfoldings are identified by red arrowheads; other myelin redundant structures (”whorls”) are indicated by blue arrows. Scale bars: 0.5 μm, except in image 2 of SC: 1 μm. **(E)** Quantification of myelin outfoldings profiles (CTR=1.30±0.22; *P2* cKO=4.15±0.55, *P* =0.0007) and myelin whorls (CTR=1.55±0.22; *P2* cKO=2.63±0.059; *P* =0.082) in ON (*n*=6). **(F)** Quantification of myelin outfoldings profiles (CTR=3.85±1.09; *P2* cKO=9.47±1.42; *P* =0.024) and myelin whorls (CTR=2.70±0.66; *P2* cKO=8.60±2.55; *P* =0.048) in SC (*n*=3/4). Statistical analysis used unpaired two-tailed Student’s *t*-test **(B, C, D,F)**. Bars represent mean±s.e.m. **(D,F)**.

### Pinch2 regulates Rho GTPase activation

We and others previously showed that myelin formation depends on tight regulation of Rho GTPase mediated signaling (Nodari et al., 2007, Pereira et al., 2009, Thurnherr et al., 2006, Benninger et al., 2007). Rho GTPases are molecular binary switches controlled by interchanging from active guanosine triphosphate (GTP)-bound state into an inactive GDP-bound form. They localize at focal adhesions (Legate et al., 2009) and are expressed in oligodendrocytes (Liang et al., 2004, Erschbamer et al., 2005) - where they are activated in a beta1-integrin-dependent mode (Feltri et al., 2008). Hence, the Ilk expression has also been shown to regulate Cdc42 and Rac1 activation in epithelial cells (Filipenko et al., 2005) and also in Schwann cells in the PNS (Pereira et al., 2009). Rho GTPases regulate downstream targets that are directly associated with the dynamics of actin cytoskeleton reorganization and local actin disassembly is fundamental in CNS myelin wrapping (Zuchero et al., 2015). We next examined the levels of classical Rho GTPases in ON lysates obtained from P30 conditional KO and control mice. First, we measured RhoA activation levels using a rhotekin-GST fusion protein to pulldown the activated form of RhoA in ON lysates. We found that the levels of the active GTP-bound form in *Pinch2* cKO ONs were significantly reduced compared with those in control ONs (**Fig. 4A,B**; 1.38±0.079 *vs* 0.62±0.095, *P* =0.0036, *n*=3), while no significant changes were observed in the total amounts of RhoA (0.96±0.07 *vs* 1.04±0.06, *P* =0.4, *n*=3). To assess the *in vivo* relevance of our observations, we analyzed myelin ultrastructure in the *Rhoa* conditional mice (*Rhoa* cKO), obtained from crossing *Rhoa*^*flox/flox*^ mice (Jackson et al., 2011) with *Cnp*^*Cre/+*^ mice (Lappe-Siefke et al., 2003) **(supplementary material Fig. S6A)**. We found that at P25 *Rhoa* cKO ON contained axons with thicker myelin when compared with controls (**Fig. 4C,D**; 0.817±0.004 *vs* 0.789±0.004, *P* =0.0096, *n*=3), suggesting that Ilk/Pinch2 signaling is required to activate RhoA and that suppression of RhoA activity elicits myelin growth.

**Figure 4.**
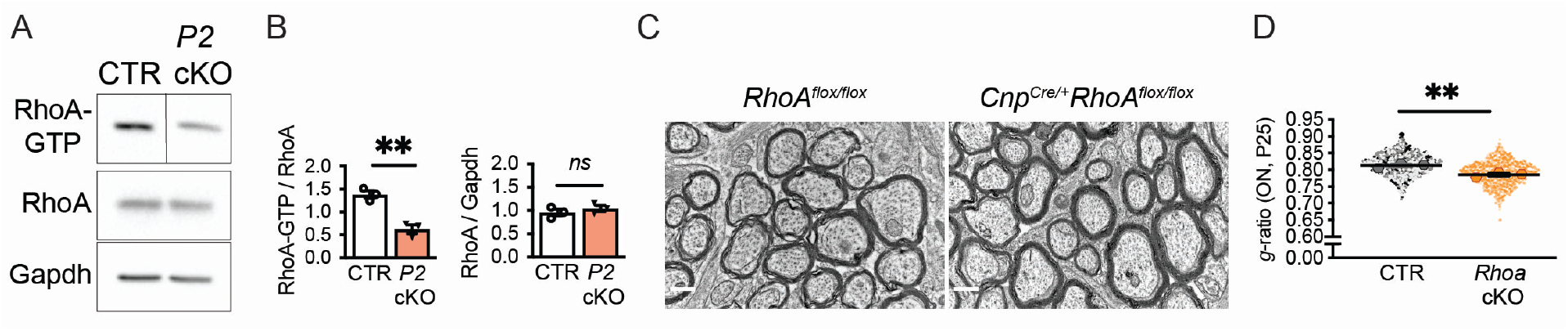
Downregulation of RhoA activity and myelin overgrowth. **(A)** Pull down assay for active GTP-bound RhoA in lysates using ON lysates from P30 *P2* cKO and control. **(B)** Quantification shows less active RhoA in *P2* cKO in comparison with control (*P* =0.004, *n*=3), and no significant changes of total RhoA expression (*n*, 2 pooled nerves). Statistical analysis used unpaired two-tailed Student’s *t*-test. **(C)** Representative EM micrographs of cross-sections of ON from conditional *Rhoa* KO (*Cnp*^*Cre/+*^ *Rhoa*^*flox/flox*^) and the respective control (*Rhoa*^*flox/flox*^) at P25. Scale bars: 0.5 um **(D)** *Rhoa* cKO displays significantly lower *g*-ratio. (*P* =0.0096, *n*=3). Beeswarm superplot shows mean *per* animal used to calculate average, SEM, and *P* value (differences with unpaired two-tailed Student’s *t*-test). Smaller symbols represent individual measures *per* animal (minimum 176 myelinated axons).

Remarkably, ablation of Cdc42 in OL results in myelin abnormalities (Thurnherr et al., 2006) similar to those found here in the *Pinch2* cKO. Next, we measured the levels of active Cdc42 in ON from *Pinch2* cKO using a glutathione-S-transferase (GST)-PAK-CD protein fusion (Sander et al., 1998), as before (Pereira et al., 2009, Benninger et al., 2007, Montani et al., 2014), and we found a significant decrease in the active form of Cdc42 (1.37±0.15 *vs* 0.73±0.10, *P* =0.025, *n*=3) in the mutant compared with the control **(Fig. 5A,B)**. In striking contrast, no significant changes were detected in *Pinch1* cKO nerves (*P* =0.20, *n*=6; **supplementary material Fig. S7A,B**).

**Figure 5.**
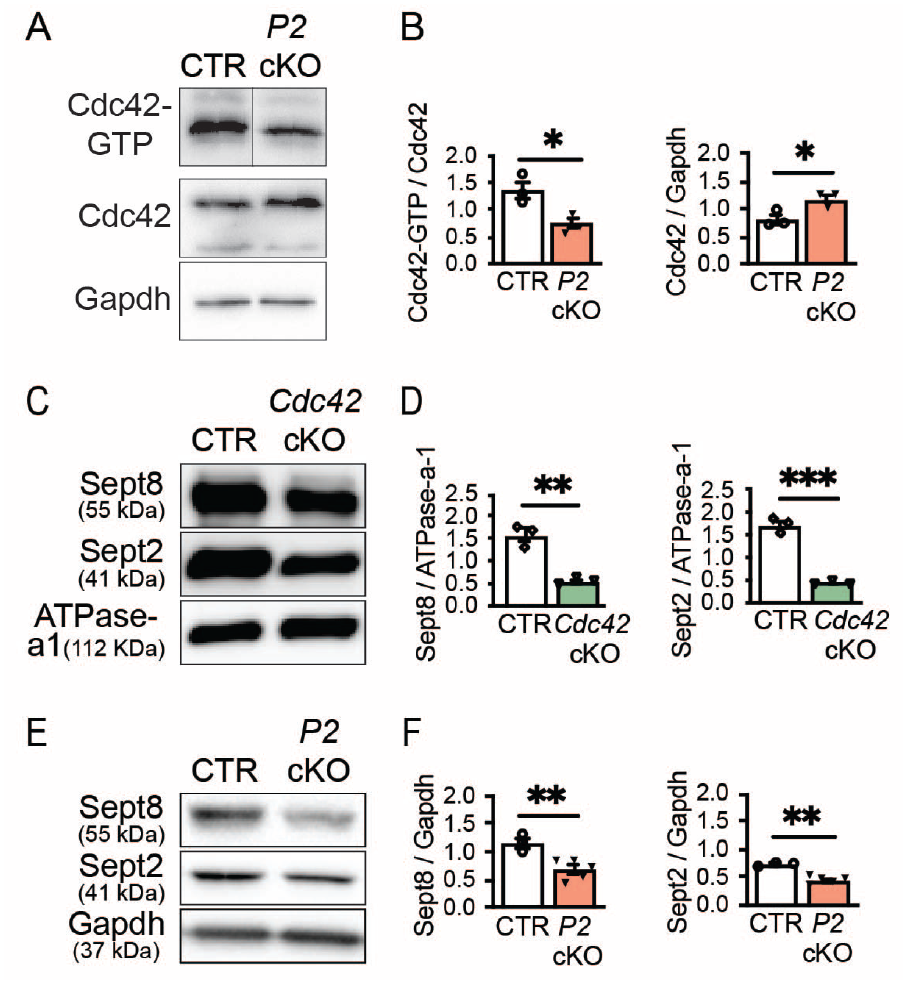
Cdc42 activation is deficient in *Pinch2* conditional knockout. **(A,B)** Pull-down assay in lysates of *P2* cKO shows decrease of the active forms of Cdc4 (*P* =0.03, *n*=3) compared with CTR (P30). Total expression of Cdc42 (*P* =0.04, *n*=3). Each *n* represents 4 pooled ONs. **(C,D)** The abundance of Septins 8 and 2 is reduced in myelin purified from brains of *Cdc42* cKO mice at P45 (Sept8: *P* =0.004; Sept2: *P* =0.008; *n*=3). ATP1A1 used as control. **(E,F)** Septins 8 and 2 are downregulated in ON from *P2* cKO (Sept8: *P* =0.009, *n*=3/5. Sept2: *P* =0.001, *n*=3/5). Statistical analysis used unpaired two-tailed Student’s *t*-test. Bars represent means±s.e.m. **(B,D,F)**

Cdc42 is involved in recruitment and oligomerization of cytoskeletal septins (Sadian et al., 2013), which form membrane-associated filaments (Bridges et al., 2014, Sirajuddin et al., 2007) whose assembly determines membrane binding and rigidity (Jiao et al., 2020, Gilden and Krummel, 2010). In myelin, septin filaments extend in the adaxonal compartment, thereby scaffolding mature myelin (Patzig et al., 2016). We found that the abundance of two of the most abundant septins in myelin, septins 2 and 8 was significantly reduced in myelin biochemically purified from the brains of *Cdc42* cKO mice compared to controls **(Fig. 5C,D)**. Indeed, OL-specific ablation of *Sept8* causes the formation of myelin outfoldings (Patzig et al., 2016) similar to those seen in *Cdc42* (Thurnherr et al., 2006) and *Pinch2* cKO. Therefore, we examined the expression of septins in protein lysates obtained from *Pinch* cKO ONs and respective controls. We found that Sept2 and Sept8 expression levels were significantly reduced in *Pinch2* cKO lysates compared with those from controls (0.73±0.02 *vs* 0.45±0.03, *P* =0.0011, *n*=3-5 (Sept2); 1.17±0.09 *vs* 0.69±0.08, *P* =0.0090, *n*=3-5 (Sept8); **Fig. 5E,F)** and were not changed in age-matched *Pinch1* cKO nerves (*P* =0.87 and *P* =0.93, *n*=3); **supplementary material Fig. S7C,D)**. Together these results suggest that Pinch2-dependent regulation of Cdc42-via control of the assembly of the septin proteins scaffold-underlie the establishment of structurally normal myelin.

## Discussion

In this study, we investigated the functions of Pinch1 and 2 during developmental myelination using oligodendrocyte (OL)-targeted gene ablation in mice. Our data show that while one isoform can functionally compensate the loss of the other during early axon ensheathment, Pinch2 is essential for subsequent myelin wrapping. We found that loss of Pinch2 leads to striking changes in myelin structure, including the formation of thicker myelin sheaths, the occurrence of myelin outfoldings, and alterations in paranodes and nodes of Ranvier. These myelin configurations emerge as the result of Pinch2 modulation of Rho GTPases as suggested from the decrease in RhoA and Cdc42 activities and from the observation that the corresponding conditional knockouts phenocopy, at least in part, the structural myelin alterations we observed in *Pinch2* cKO. Thus, our study supports the view of a bimodular arrangement of Pinch isoforms in IPP as a defining aspect of OL function along development and shows for the first time that Pinch2 coordinates essential Rho GTP signaling in myelination **(Fig. 6)**.

**Figure 6.**
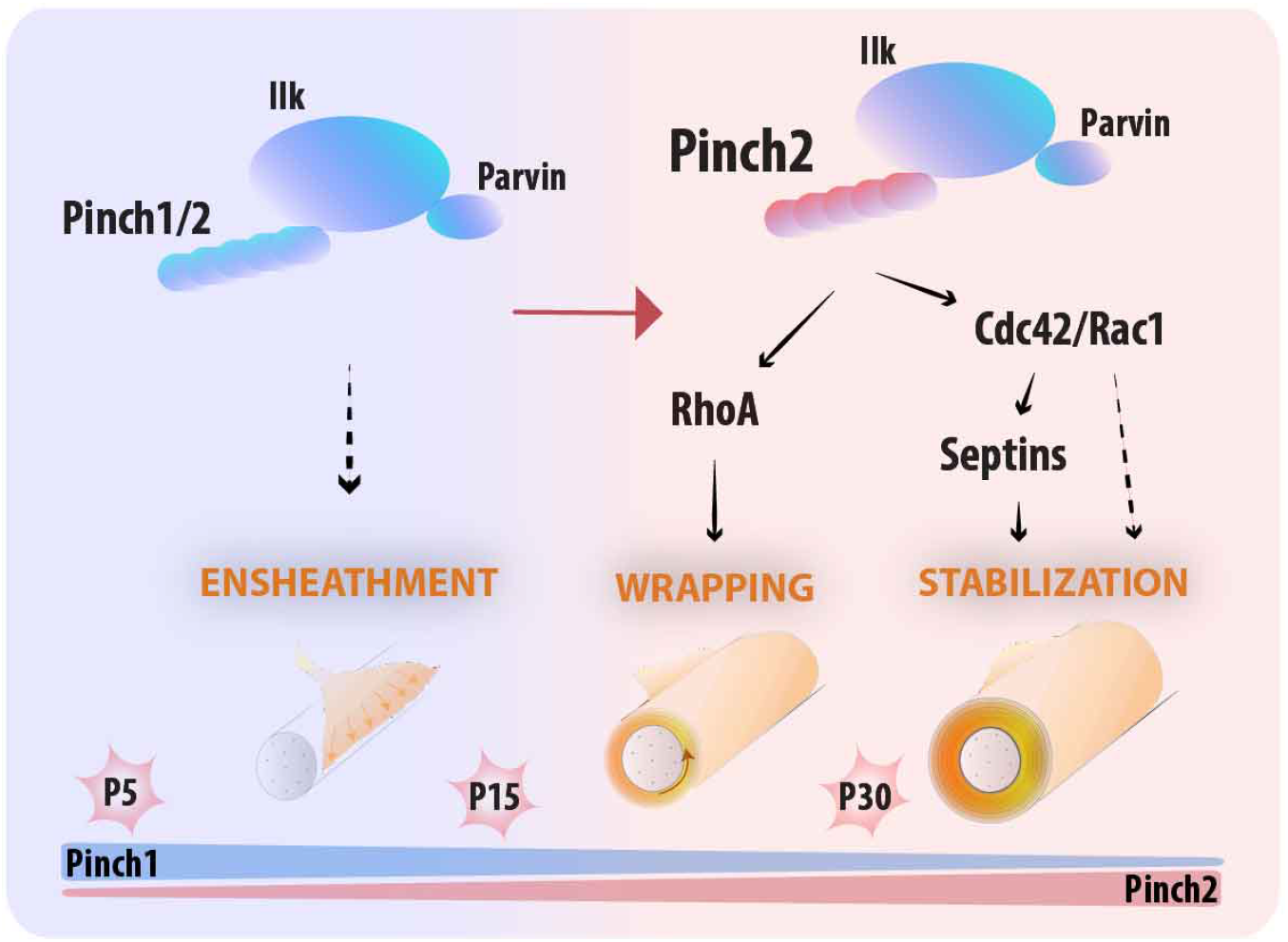
Ilk-Parvin-Pinch (IPP)-mediated signaling is required for the correct formation of myelin sheaths in the CNS. Axon contact and ensheathment occurs in the presence of either Pinch isoform, but Pinch2 specifically controls myelin thickness and stability by modulating the activities of RhoA, Cdc42 and Rac1 Rho GTPases in the myelinating OL.

The possibility that the two Pinch proteins have partially redundant functions between them has been suggested before (Fukuda et al., 2003, Xu et al., 2005) but has never been formally demonstrated *in vivo*. Similar to other cell types, Pinch1 expression is detected earlier than Pinch2 in the developing OL. We found that the recruitment of either Pinch supports the integrity of IPP complex during the initial stages of axon ensheathment and early myelination. Upon *Pinch1* loss and despite the absence of an obvious myelin phenotype, there is a transient increment of Olig2-positive cells along with an increase in the expression of the extracellular signal-regulated protein kinase (Erk) 1 and 2, a known regulator of OPC proliferation (Ishii et al., 2013). The increase in OPC number is balanced by augmented OL apoptosis likely through the transient activation of Jnk **(supplementary material Fig. S2)** (Kadrmas et al., 2004), and is in line with the previously described pro-survival role of Pinch1 in the primitive endoderm (Montanez et al., 2012). This function of Pinch1 may rely on the binding to the Ras suppressor (Rsu1) protein, a negative regulator of Jnk signaling (Kadrmas et al., 2004) expressed in OLs (Zhang et al., 2014). Globally, this supports the involvement of Pinch1 in adjusting myelinating glia homeostasis during development. This capacity of the developing nervous system to counteract the initial excess of OLs (Raff et al., 1998), is thought to require integrin-mediated signaling (Benninger et al., 2006, Colognato et al., 2002, Laursen et al., 2009). During development Pinch2 protein expression is detected and peaks later than Pinch1 protein that declines after the second postnatal week, when myelination in the optic nerve progresses more intensely (Dangata and Kaufman, 1997, Foran and Peterson, 1992). At this stage, we observed a significant and progressive change in myelin structure (increase in the thickness of myelin sheath and in the emergence of myelin outfoldings) in *Pinch2* cKO mice. Thus, the developmentally-driven increase in the abundance of Pinch2 and consequent change in the molecular composition of the IPP complex, - IP(Pinch2)-appears essential for myelin formation and maturation. Along with the obligatory binding partners Parvin and Ilk, Pinch form the IPP protein complex that bridges integrins and the actin cytoskeleton (Wickstrom et al., 2010, Legate et al., 2006). The tripartite IPP associates with various actin-binding proteins, including kindlin2 (Bledzka et al., 2016), paxillin and thymosin-B4 (Bock-Marquette et al., 2004) and Ilk protein determines F-actin bundling through coordinated F-actin binding of actin to Pinch1 and Parvin (Vaynberg et al., 2018). Depletion of *a-Parvin* (Montanez et al., 2009) or *Ilk* (Kogata et al., 2009) in mice vascular smooth muscle cells (vSMCs) impair proper vessel formation by RhoA-mediated changes in actomyosin contractility. In the peripheral nervous system, loss of *Ilk* in Schwann cells results in upregulation of Rho/ROCK signaling and hampers the extension of cytoplasmic processes arresting radial sorting and, as a consequence, myelination (Pereira et al., 2009). Ilk regulates actin cytoskeleton dynamics and morphogenesis in OLs (O’Meara et al., 2013, Michalski et al., 2016). RhoA-elicited morphological changes and OL development have been studied *in vitro* (Liang et al., 2004, Wang et al., 2012, Rajasekharan et al., 2010) and support the view that OL process extension depends on the precise temporal modulation of active RhoA. Although we showed that actin filament assembly is initially required to form myelin segments in myelinating co-cultures (Azevedo et al., 2018), studies by Pedraza et al. indicated that inhibition of the RhoA downstream effector Rho kinase (ROCK) leads to enhanced formation of myelin segments in coculture (Pedraza et al., 2014). This is in line with the observation that OL membrane spreading and myelin wrapping *in vivo* requires the loss of actin filaments as a result of the action of actin severing and actin depolymerization proteins (Zuchero et al., 2015, Nawaz et al., 2015). In the present work, we found a significant increase in the number of myelin wraps upon *Pinch2* depletion associated with a decrease in RhoA activity, suggesting that RhoA may be negatively controlling myelin sheath growth and preventing overmyelination. As in *Pinch2* cKO nerves PI3K-AKT and PTEN signaling, known to drive myelin membrane growth (Harrington et al., 2010, Goebbels et al., 2010, Flores et al., 2008, Ishii et al., 2013) were not significantly affected, reduced RhoA activity could result in inhibition of actin nucleation via the Rho GTPase effectors Diaphonous-related formins (Kuhn and Geyer, 2014), ultimately leading to decrease of actin filaments and augmented membrane expansion and myelin wrapping. Of note, actin dynamics can also be controlled by Rsu1 protein (Kadrmas et al., 2004, Gonzalez-Nieves et al., 2013, Yang et al., 2021) and Rsu1 transcript is reduced along OL development (Zhang et al., 2014) and in *Pinch2* cKOs (*our results*).

We and others have shown that cytoskeleton dynamics regulate different aspects of OL biology, including axonal wrapping and the proper formation of myelin sheaths (reviewed in Hughes and Appel, 2016, Domingues et al., 2018, Brown and Macklin, 2020). Analysis of the *Pinch2* cKO nerves also showed defects in paranodal axoglial junctions at the end of each internode **(supplementary material Fig. S5F)** and in the length of the node of Ranvier (NR) **(supplementary material Fig. S5G–I)**. Changes in the length of the NR may reflect an attempt to reestablish normal action potential conductance speed in response to myelin overgrowth. Increasing the length of the node could change capacitance and increase conduction speed along the nerve, potentially compensating for alterations in conduction along over myelinated axons. This is in line with the notion that NR length is a regulator of myelinated axon conductance in the CNS (Arancibia-Carcamo et al., 2017). Flanking the nodes, the integrity of paranodes relies on integrin signaling (Laursen et al., 2009) and on axon-glia adhesions (Southwood et al., 2004) that once dysregulated could also result in abnormally thick myelin sheaths (Elazar et al., 2019).

Myelin outfoldings occur during normal development (Patzig et al., 2016, Snaidero et al., 2014) and are also a hallmark of age-dependent pathology in the CNS (Peters, 2002). In the mature CNS, an increased frequency of myelin outfoldings has been observed upon experimentally increasing PIP3 levels (Goebbels et al., 2010), changing actin organization and dynamics (Zuchero et al., 2015, Katanov et al., 2020) and septin assembly (Patzig et al., 2016; Erwig et al., 2019). *Pinch2* cKO-in which we observed a reduction in the pool of active Cdc42 (and Rac1, not shown)-displayed myelin outfoldings similar to those previously reported for OL-*Cdc42* /*Rac1* conditional knockouts (Thurnherr et al., 2006). Importantly, *Cdc42* /*Rac1* cKOs do not exhibit thicker myelin sheaths, which further supports the existence an independent pathway limiting myelin wraps in *Pinch2* cKO.

Cdc42 has an essential role in the regulation of assembly/ disassembly of the septin complex, and the introduction of specific mutations in the budding yeast *Cdc42* gene that results in defective basal Cdc42 GTPase activity, leads to defective septin assembly (Gladfelter et al., 2002). This is consistent with our data showing reduced septin expression in *Cdc42* cKO ONs. Cdc42 GTPase negatively regulates the Borg proteins that control the organization of septin network complex (Joberty et al., 2001). For example, Borg5/Cdc42ep1/Cep1 is an effector of Cdc42 (Bagci et al., 2020) expressed in OLs (Zhang et al., 2014) and identified in myelin (Jahn et al., 2020). Filamentous septins were identified in the innermost layer of the myelin sheath assembled as longitudinal filaments in the non-compact adaxonal region of the membrane (Patzig et al., 2016). The assembly of the myelin septin filament correlates with myelin sheath stability, as suggested by the formation of myelin outfoldings when septin 8 is depleted (Patzig et al., 2016). Two subunits of the core myelin septin oligomer, Septins 2 and 8, are significantly reduced in *Pinch2* cKO myelin. Because oligomer formation depends on the availability of individual septins (Hall et al., 2005), the decrease in Sept 2 and 8 likely elicits the assembly of fewer oligomers and underlies the formation of myelin outfoldings in *Pinch2* cKO similar to those found in *Sept8* cKO (Patzig et al., 2016).

We note that *Lims2* /*Pinch2* expression is increased during aging in the oligodendrocyte lineage (Ximerakis et al., 2019), which suggests a critical role of Pinch2-mediated signaling with age. In aged non-diseased brains, the occurrence of myelin outfoldings (Peters, 2002, Patzig et al., 2016), similarly to those observed in *Pinch2* cKO (and *Cdc42* cKO (Thurnherr et al., 2006)), positively correlate with the age-associated decline in neurological function (Gunning-Dixon and Raz, 2000, Peters et al., 1996, Liu et al., 2017) and could reflect an increasingly less compliant cytoskeletal structure (Seixas et al., 2019) to maintain myelin stability. Hence, future studies on the IPP-mediated molecular mechanisms underlying the stabilization of myelin sheaths in the aging CNS could be particularly relevant in the comprehension of age-associated systemic brain changes and cognitive impairment.

## Materials and Methods

### Animals

All experiments involving animals were subjected to prior approval by the local Ethics Committee and were conducted in strict compliance with European and Portuguese guidelines (Project Licences DGAV 11770/2014 and 002803/2021). *Mus musculus* (mice) were co-housed in groups up to 6 animals, fed with standard rodent diet and water *ad libitum* and kept in a 12 hours light /dark cycle. Mice with *floxed* alleles for *Lims1* (Li et al., 2005) (*Pinch1*^*flox/flox*^ or *Pinch1*^*flox/wt*^), *Lims2* (Stanchi et al., 2005) (*Pinch2*^*flox/flox*^ or *Pinch2*^*flox/wt*^), *Rhoa* (Jackson et al., 2011) *Rhoa*^*flox/flox*^ or *Rhoa*^*flox/wt*^ and *Cdc42* (Wu et al., 2006) (*Cdc42*^*flox/flox*^ or *Cdc42*^*flox/wt*^) have been described previously. For oligodendroglia-specific conditional depletion of *Lims1* /*Pinch1*, *Lims2* /*Pinch2*, *Cdc42*, and *Rhoa* mice carrying *floxed* alleles were crossed with the *Cnp*-*Cre* driver line (Lappe-Siefke et al., 2003). In all lines, experimental mice were on C57BL/6 background. In the offspring the relevant genotypes were determined by genomic PCR using the following primers: Pinch1 fw 5’-CTA GGC TGG TAA TGC AGG CC-3’ and rv 5’-CCT GCC AAT GAT GAA TTC AC-3’ [expected PCR product 230 bp (wt) and 400 bp (floxed)]; Pinch2 fw: 5’-CAC TCC CAA TTC CCC TCC CTG AG-3’ and rv: 5’-AGG GGT CTG AGG TCC TGA GAA GG-3’ [expected PCR product 298 bp (wt) and 438bp (floxed)]; Cdc42 fw: 5’-TCT GCC CTG ATC TAC ACA TAC AC-3’ and rv: 5’-ATG TAG TGT CTG TCC ATT GG-3’ [expected PCR product 200 bp (wt) and 300 bp (floxed)]. RhoA fw: 5’ - AGC CAG CCT CTT GAC CGA TTT A - 3’ and rv 5’ - TGT GGG ATA CCG TTT GAG CAT - 3’ [expected PCR product 297bp (wt) and 393 bp (floxed)] - 3’. Cnp fw: 5’-GAT GGG GCT TAC TCT TGC-3’ and rv: 5’-CAT AGC CTG AAG AAC GAG A-3’ (expected PCR product 900bp). Cre fw: 5’-ACC AGG TTC GTT CAC TCA TGG-3’ and rv: 5’-AGG CTA AGT GCC TTC TCT ACA-3’ (expected PCR product 230 bp). Conditional knock-out (cKO) genotypes are referred to as follows *Pinch1* cKO, *Pinch2* cKO, *Rhoa* cKO and *Cdc42* cKO. In all experimental setups, aged-matched males and females were arbitrarily used.

### Transcardial perfusion and tissue fixation

Adult mice were anesthetized by terminal intraperitoneal administration of pentobarbital (0.2 ml/ 30 g body weight) and subsequently transcardially perfused with sterile, filtered and pre-warmed at 37°C, phosphate buffer saline (PBS) or 0.1 M phosphate buffer (PB) pH 7.4, depending if the dissection of specimens was for regular histology or to be used in electron microscopy (EM). This was immediately followed by pre-warmed (37°C) fixative perfusion, using 4% paraformaldehyde solution prepared in PBS (samples for immunocytochemistry, IHC), or 4% paraformaldehyde/ 3% glutaraldehyde solution prepared in PB (samples for EM). After dissection, the tissue of interest for IHC was incubated overnight (O/N) in 4% PFA/PBS, and then immersed in sterile 20% (w/v) sucrose/PBS until tissue reached the bottom of the reservoir for optimal tissue infiltration. Fixed samples were embedded in OCT (Criomatrix), and stored at −80°C. Samples used in EM were post fixed at least O/N in the fixative solution.

### Transmission electron microscopy

White matter samples were visualized using transmission electron microscopy (TEM) in perfusion fixed tissue. Postnatal days 15 (P15), young adult (P30) or adult at P90 were fixed by cardiac perfusion with 4% paraformaldehyde/ 3% glutaraldehyde in 0.1 M phosphate buffer, pH 7.4. After dissection, the relevant regions from central nervous system (optic nerve, thoracic spinal cord, corpus callosum) were left in perfusing buffer for at least 12 hours, and then post fixed overnight in 1% Osmium Tetroxide in 0.1 M phosphate buffer. Samples were then dehydrated through increasing ethanol concentrations and finally resin-embedded. Control and KO material was handled in parallel to avoid misinterpretation of myelin changes and the discard of myelin artifacts. Blocks were processed in a ultramicrotome to obtain 1 μm thin (Thin sections) that were stained with toluidine blue to assess the integrity of samples and to select further the tissue for analysis. Next, 60 nm sections (Ultrathin sections) from the selected areas were cut, counterstained using 3% uranyl acetate and 1% lead citrate, transferred onto slotted grids and visualized using the electron microscope (Jeol JEM 1400) at 80 Kv, coupled to an Orius Gatan camera. Standard hexagonal mesh grids were used, except when larger samples were imaged for quantification purposes. In this case, were used single hole grids coated with pioloform (0.7% in chloroform solution). At least 3 animals per genotype were visualized. To obtain representative high-resolution images, ultrathin sections were imaged, typically at 25,000x, 12,000x and 30,000x for optic nerve, spinal cord and corpus callosum, respectively. For quantifications, images were taken at 12,000x. Image processing was performed using ImageJ 1.52h (Schindelin et al., 2012) and Photoshop CS5.

### Quantification and morphometric analysis

Both males and females were used in the study. All the quantifications were performed with the researcher(s) blind to the genotypes of the mice. For EM, randomly digitalized and non-overlapping images of the optic nerve or ventral spinal cord were analyzed for *g*-ratios by measuring at least 100 axons *per* animal. In order to measure contacted axons (P15), 6% overlapping images from half of the ON were stitched (Photomerge, Photoshop). For western blots, the results were compared solely among the same membrane blot and if possible using animals from the same litter (the rare cases where this was not possible, are stated in the text). Bands were quantified using ImageLab, except in the case of blots resulting from pull-downs, in which, comparison involves samples from different membranes. In either case, blots displaying areas of the bands with saturated signal intensity (above a measurable range at ImageLab) were discarded. For morphological analysis (*g*-ratios) on ultrathin sections, at least 10 images *per* animal were randomly taken at 12,000. *G*-ratios were calculated by dividing the area of the axon, inferred from the measured perimeter, by the area of fiber (axon with myelin), also inferred from the measured perimeter. At least 100 axons were measured from at least 3 animals *per* genotype. At P30, we used an alternative method in parallel, so that the diameter of compacted myelin (replacing axon diameter), was divided by the outer diameter of myelin sheath (Goebbels et al., 2010). To determine the number of contacted/myelinated axons in the optic nerve, quantification was carried after image alignment (Photoshop CS6) from an hemi section of the optic nerve (P15 days). To quantify myelin outfoldings profiles: 741-1000 **(Fig. 3E)**, 397-849 **(Fig. 3F)**, 1600 **(Fig. S5B)**, 450 **(Fig. S5C)**, 320 **(Fig. S5E)** were examined in at least 10 random and non-overlapping frames *per* animal (12K EM images). To quantify the length of nodes of Ranvier and the diameter of nodes and axons at node **(Fig. S5J–M)**, between 75-87 axons of SC and 47-81 axons of ON *per* genotype were examined. Every quantification involving EM images was manually made using ImageJ.

### Western Blot

For protein analysis, the indicated region of central nervous system was dissected and immediately frozen. The frozen tissue was homogenized using a chilled micropestle in cold RIPA buffer (Sigma, R0278), supplemented with 1 mM EDTA and protease and phosphatase inhibitors, according to the manufacturer’s instructions. Samples were cleared in pre-cooled centrifuge for 15 minutes at 4°C under maximum speed. For optic tissue, and unless stated, two nerves from one animal were loaded per lane. For spinal cord, protein concentration was measured using the colorimetric BCA Protein assay. 10 μg of protein *per* lane were resolved in 10% or 12% SDS-polyacrylamide gels, after dilution in GLB (150 mM Trizma Base, 6% SDS, 0.05% bromophenol blue, 30% glycerol, and 6 mM EDTA pH 8.8) and freshly added DTT (1 mM). Heat-induced denaturation of proteins was achieved by ten minutes at 95°C immediately before loading. The pre-stained marker Precision Plus Protein Dual Color Standard (Biorad) was used to determine approximate molecular weights and monitor transfer efficiency. Before blocking, protein bands were visualized by Ponceau S staining. For the immunoblots displayed in **Fig. 5E**, myelin was purified from the brains of mice as previously reported (Patzig et al., 2016) and separated on 10% gels with a protein load of 5 μg *per* lane (SEPT2) or 10 μg *per* lane (SEPT8, ATP1A1). Instead of heat denaturation, samples were incubated 10 minutes at 42°C before loading. Nitrocellulose membranes were blocked for 1 hour in 5% nonfat dry milk in TBS-Tween (TBS-T) or 90 minutes in 5% BSA-TBS-T. Incubation with primary antibodies in blocking solution overnight, with rotation at 4°C, was followed by membrane washing with TBS-T for 30 minutes and incubation with HRP-conjugated secondary antibodies at room temperature. After washing, blots were incubated with Super Signal West Pico Chemiluminescent Substrate (Fisher) reagent for band detection and the signal was detected using ChemiDoc Imaging System (XRS, Bio-Rad) coupled to ImageLab software.

### Immunostaining

Paraformaldehyde (4% PFA)-fixed samples of *Pinch1* cKO or *Pinch2* cKO and the respective controls were cut in 14 um thin cross sections using a cryostat (Leica CM 3050S). KOs and the respective control animals were always transferred to the same slide (SuperFrost Plus coated slides). To minimize bias in processing the tissues, 5-12 sections *per* animal were transferred to a slide, and simultaneously processed before blinded subjected to analysis. For fluorescence staining, sections were rinsed in TBS for 10 minutes, permeabilized with 0.5% Triton X-100 in TBS for 15 minutes and blocked for 1 hour in blocking solution (10% normal goat serum, 1% BSA, 0.025% Triton X-100 in TBS). Samples were then incubated over night at 4°C in 1% BSA and 0.025% Triton X-100 in TBS. After washing three times with 0.025% Triton X-100 in TBS (TBS-T) for 15 minutes, samples were incubated with the secondary antibody for at least 1 hour in TBS with 1% BSA at room temperature. The samples in the slide were washed 3 times with TBS-T, interleaved with the incubation with 4,6-diamidino-2-phenylindole (DAPI, Alfagene) for 10 minutes. Slides were mounted using Fluoroshield (Sigma Aldrich) and left to dry prior to imaging.

### Antibodies

Antibodies against Pinch proteins were raised using the C-terminal based peptides for immunization: CLKKLSETLGRK for Pinch1 and CAQPKSVDVNSL for Pinch2 (Li et al., 2005). The antibodies detected Pinch1 or Pinch2 at the estimated apparent molecular weights, using a 1:1000 dilution under standard denaturing conditions. Other primary antibodies used in western blot analysis were: Ilk (BD Biosciences, 611803, 1:1000), a-Parvin (Abcam, #11336, 1:3000), Akt (Cell Signaling, #9272, 1:1000), p-Akt Thr 308 (Cell Signaling, #2965, 1:1000), p38 (Cell Signaling, #8690, 1:1000), p-p38 (Cell Signaling, #4511, 1:1000), Erk (Cell Signaling, #9102, 1:1000), p-Erk (Cell Signaling, #9101, 1:1000), Jnk (Cell Signaling, #9258, 1:1000), p-Jnk (Cell Signaling, #4668, 1:1000), Pten (Cell Signaling, #9188, 1:1000), Sept8 (ProteinTech, #11769-1-AP, 1:2500), Sept2 (ProteinTech, #11397-1-AP, 1:500), ATP1A1 (Abcam, #ab7671, 1:2500), Cdc42 (Cell Signaling, #2466, 1:1000), RhoA (67B9) (Cell Signalling, #2117, 1:1000), Gapdh (HyTest, #6C5, 1:50,000), Tubulin (Sigma, #T5168, 1:10,000). Either Gapdh or Tubulin were used as reference to avoid superimposed bands in blots. All these antibodies were used in standard conditions (blocking and antibodies incubation in 5% low fat milk in TBS-T), except the antibodies for Cdc42 and for the phosphorylated forms of the proteins, which were incubated in 5% BSA in TBS-T. In immunostaining the following primary antibodies were used: Olig2 (Millipore, #AB9610, 1:500), CC1 (Calbiochem, OP80, 1:200). Peroxidase-Conjugated AffiniPure Donkey Anti-Rabbit IgG (Jackson ImmunoResearch, #711-035-152. 1:10000), Peroxidase-Conjugates AffiniPure Goat Anti-Mouse IgG (Jackson ImmunoResearch, #115-035-146, 1:15000), Peroxidase-Conjugated AffiniPure Goat Anti-Rat IgG (Jackson ImmunoResearch, #112-035-003, 1:10000) were used as secondary antibodies for western blot. As for immunostaining, Alexa Fluor Anti-Rabbit 488 (Alfagene, A11008, 1.1000) and Cy 3-Conjugated AffiniPure Goat Anti-Mouse IgG (Jackson ImmunoResearch, #115-165-146, 1:1000).

### Rho Gtpase activity assay

RhoA and Cdc42 activities were measured using either a glutathione-S-transferase (GST)-rhotekin or PAK-CD (PAK-CRIB domain)-based assay, as described previously (Sander et al., 1998, Manser et al., 1994). Briefly, expression of recombinant protein GST-Rhotekin or GST-PAK-CD was induced in transformed BL21 Escherichia coli by addiction of 0.1 M isopropylthiogalactoside for 4 hours. After harvesting, bacteria were resuspended in lysis buffer (50 mM Tris-HCl, pH 8.2 mM MgCl2, 0.2 mM Na2S2O, 10% glycerol, 20% sucrose, 2 mM dithiothreitol, 1 g/ml leupeptin, 1 g/ml pepstatin, and 1 g/ml aprotinin) and sonicated at 4°C. Cell lysates were centrifuged for 20 minutes at 4°C (45,000 g) and the cleared supernatant stored at −80°C. For bait immobilization, 300 μl of supernatant was incubated with 40 μl of Glutathione High Capacity magnetic agarose beads (Sigma) for at least 30 minutes, at 4°C, with gentle agitation. Beads were washed twice using lysis buffer and twice with FISH buffer (10% glycerol, 50 mM Tris-HCl, pH 7.4, 100 mM NaCl, 1% NP-40, 2 mM MgCl2). A second tube was subjected to the same steps containing equivalent volumes of beads and FISH buffer (instead of bait). The contents of the two tubes (bait-immobilized beads and “clean” beads) was mixed and then evenly distributed in two tubes with lysates. To prepare the lysates, the ONs from at least two animals (P30) were pooled, homogenized in cold FISH buffer (with protease inhibitor cocktail) and centrifuged for 15 minutes at 4°C (13,000 g). Exactly 10% of volume was taken and frozen (input). The remaining supernatant was incubated with the bacterially produced GST-rhotekin or GST-PAK-CD fusion bound to GST-coupled magnetic beads for about 12 hours, with gently agitation at 4°C, and washed 4 times with excess of FISH buffer with cocktail of protease inhibitors. For elution, GLB (150 mM Trizma Base, 6% SDS, 0.05% Bromophenol Blue, 30% glycerol and 6 nM EDTA pH 8.8) was added and samples incubated at 95°C for 10 minutes. Samples were resolved on a 12% SDS PAGE gel (pull down samples run between empty lanes), followed by standard western blot and Ponceau S staining to confirm uniform pool downs prior detection with the relevant antibodies.

### Statistical analysis

Assumptions on normality were made based on prior testing of Gaussian distribution using Shapiro-Wilk test. Assumption of homogeneity of variances were assumed to be identical, but whenever was not met (F-Test for Equality of two variances, *P* greater than 0.05), the Student’s *t*-test was replaced by the non-parametric Mann-Whitney U test to compare two independent groups on a continuous outcome. Statistical analyses were completed using GraphPad Prism version 8.3.1 (RRID: SCR002798) for Mac OS. Unless otherwise stated and in cases when only two conditions or samples (*e.g.* genotypes) were compared, data were analysed by unpaired two-tailed Student’s *t*-test. Multiple group analysis was performed using one-way ANOVA and the indicated *post hoc* tests in figure’s legends. Further analyses are indicated in the legends of the figures. All controls and KO mice were littermates, and often sample size corresponds to the litter size and to the numbers generally employed in the field, although no statistical method was employed for sample size determination. Outliers were not excluded and all data are expressed as “mean” and the “standard error of the mean” (SEM) to describe the variability within at least three independent samples. Scatter SuperPlots (Lord et al., 2020) were specifically designed to convey experimental variability, in which the full dataset of technical replicates was overlay with a second plot with the correspondent averages of biological replicates (*e.g*. in **Fig. 2C, Fig. 3B, Fig. 4D**).

## Data availability

Raw data available online.

## Supplementary information

The manuscript contains 7 supplementary figures.

## Acknowledgments

We thank K.-A. Nave (Max Planck Institute of Experimental Medicine, Göttingen) for the Cnp-Cre transgenic line and M. S. Erwig (Max Planck Institute of Experimental Medicine, Göttingen) for technical assistance. We acknowledge Ana Seixas (i3S, Porto) for insightful observations and critic revision of the manuscript, Jorge A. Pereira (ETH, Switzerland) and Huiliang Li (UCL, UK) for helpful discussions and critic revision of the manuscript. We thank technical support from i3S scientific platforms Histology and Electron Microscopy and Advanced Light Microscopy (members of the national infrastructure PPBI-Portuguese Platform of BioImaging, supported by POCI-01-0145-FEDER-022122), and the Cell Culture and Genotyping and the Animal Facilities.

## Contribution

Conceptualization: J.P.F., J.B.R.; Methodology: J.P.F., R.V-S., H.B.W.; Validation: J.P.F.; Formal analysis: J.P.F., R.V-S.; Investigation: J.P.F., R.V-S.; Data curation: J.P.F.; Writing - original draft: J.P.F.; Writing - review editing: J.P.F., R.F., H.B.W., J.B.R.; Visualization: J.P.F.; Supervision: J.P.F., J.B.R.; Project administration: J.P.F., J.B.R.; Funding acquisition: J.B.R.

---

**Fig. S1.**
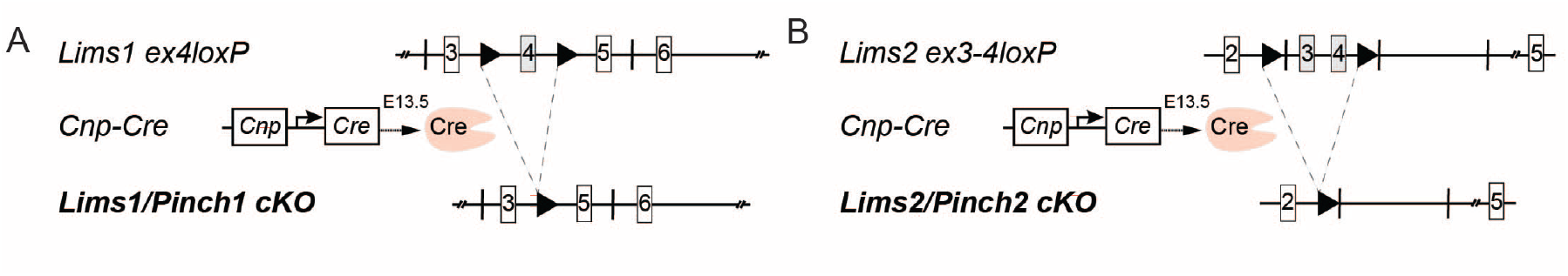
Strategy of conditional inactivation of *LimsVPinchl* and *Lims2/Pinch2* alleles in the oligodendrocyte lineage upon *Cnp*-driven *Cre* expression *in vivo*. (**A,B**) In *Pinch1*^*flox/flox*^ mice, exon 4 is flanked by *LoxP* sites (▶), whereas exons 3 and 4 are flanked in *Pinch2*^*flox/flox*^ mice. Excision occurs upon expression of *Cre* recombinase under the control of *Cnpase* gene regulatory elements.

**Fig. S2.**
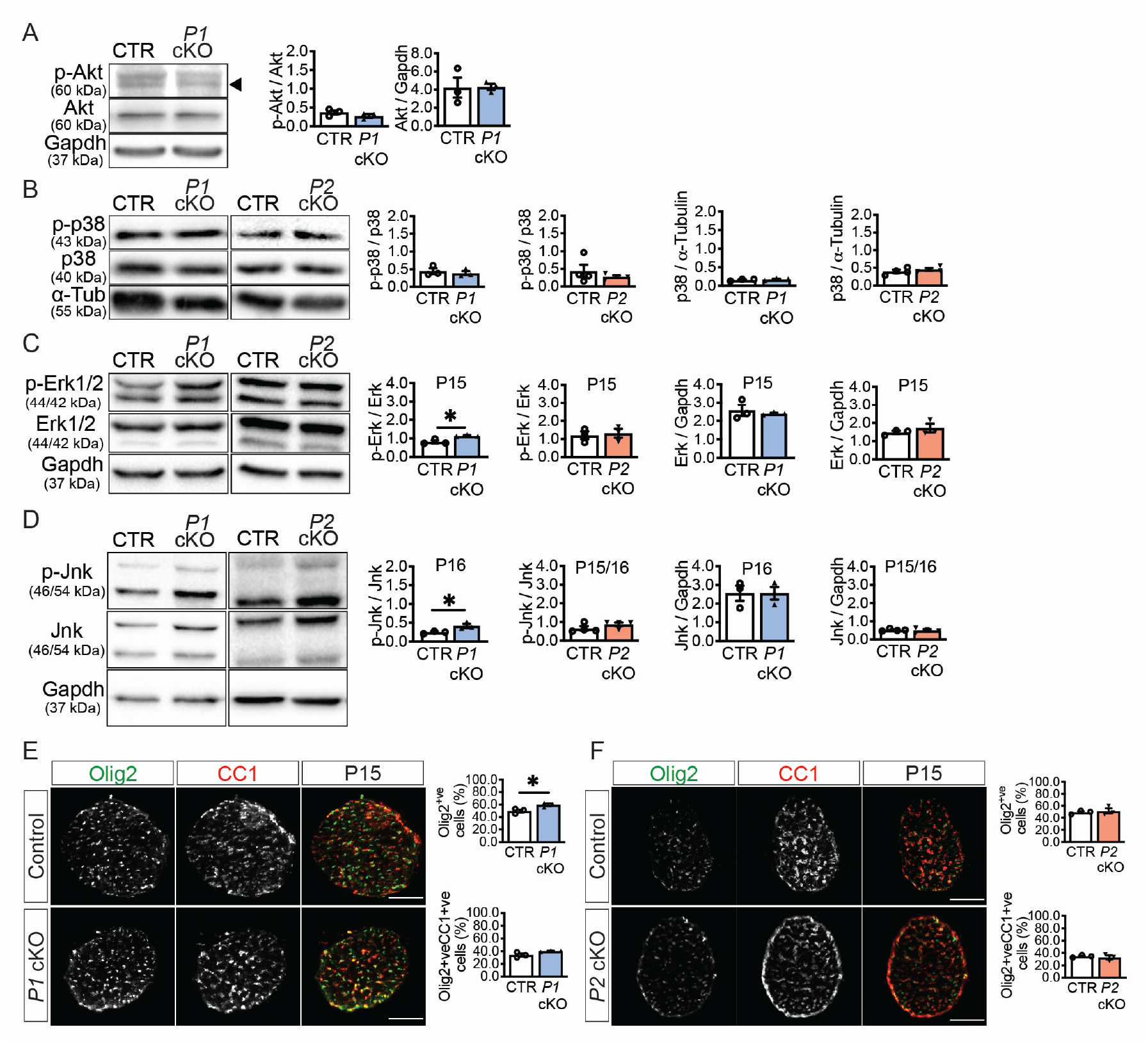
Pinchl mediates control of OL/OPC population through MAPKs. **(A,D)** Western blot analyses of ON homogenates derived from P15/16 CTR and *Pinch* cKOs, show no significant variations in phospho-Akt (Thr308) and p-38, and an increased in p-Erk1/2 (*P*=0.018, *n*=3) and p-Jnk (*P*=0.039, *n*=3) in *P1* cKO. No variation was detected in *P2* cKO. Statistical differences determined by unpaired two-tailed Student’s *t*-test (*n*=3/4), except when Fisher’s *P*<0.05 in p-p38 (*P2* cKO) and in Erk 1/2 (*P1* cKO) where Mann-Whitney test was used instead.) **(E-F)** Immunolabeling of ON cross sections of *Pinch* cKO at P15 show an increase in Olig2-positive cells in *P1* cKO (*P*=0.037), and no variation in the detected CC1-positive cells. The number of Olig2 and CC1-positice cells is similar in *P2* cKO and CTR. Statistical differences determined by unpaired two-tailed Student’s *t*-test (*n*=3). Scale bars 100 μm. Bars represent means±s.e.m. **(A-F).**

**Fig. S3.**
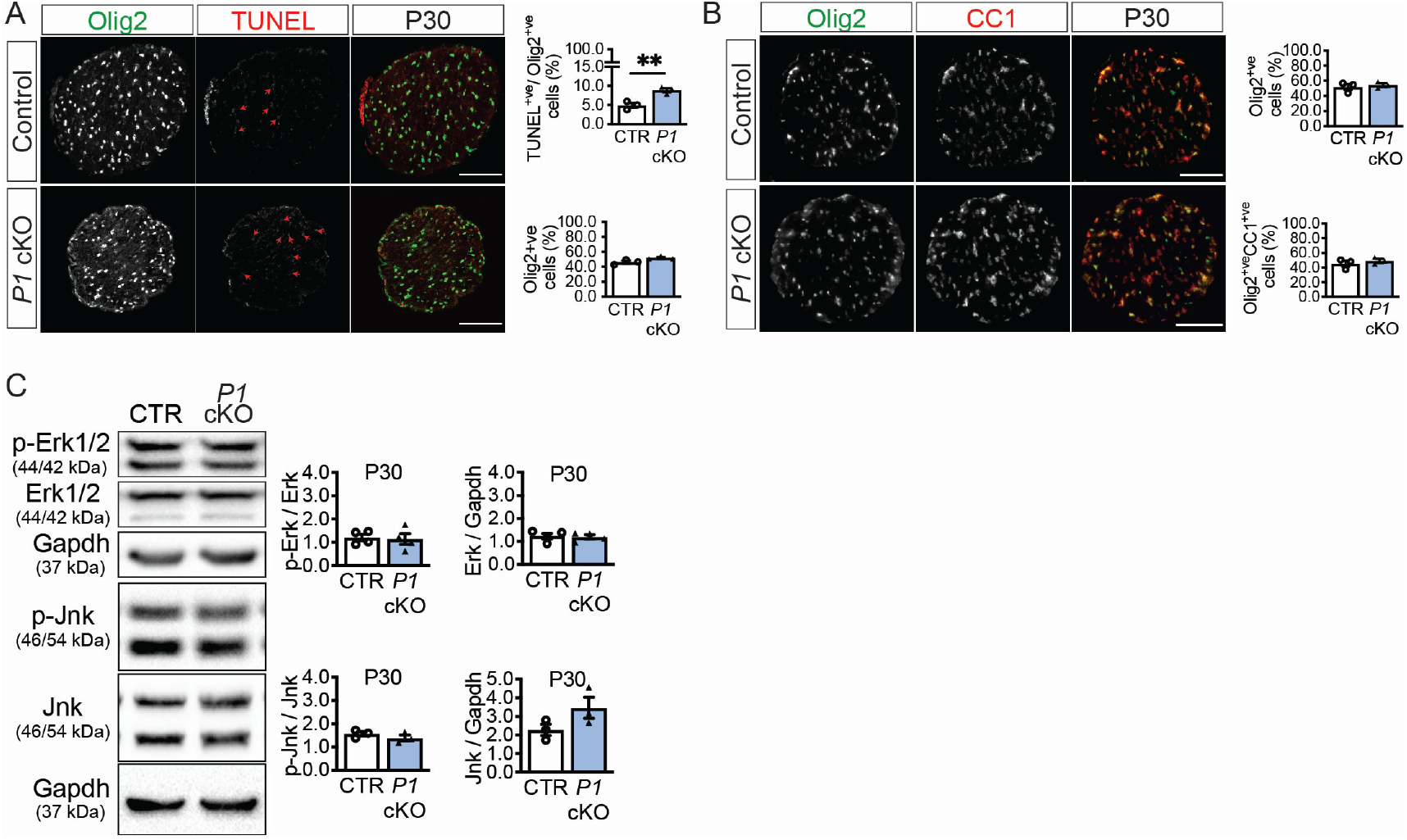
Pinchl is required to maintain OL homeostasis. **(A)** Increase in the number of TUNEL-positive cells in ON cross sections of *P1* cKO at P30 (*P*=0.009, *n*=3). Arrowheads highlight examples of TUNEL-positive nuclei. **(ET** Similar numbers of Olig2 and CC1-positive cells in ON cross sections of P1 cKO and CTR at P30. Scale bars 100 μm **(A,B). (C)** Western blot analyzes show no variation in levels of phospho-Erk and phospho-Jnk in the ON of *P1* cKO at P30. Statistical differences determined by unpaired two-tailed Student’s *t*-test (*n*=3). Bars represent means±s.e.m. **(A,B,C).**

**Fig. S4.**
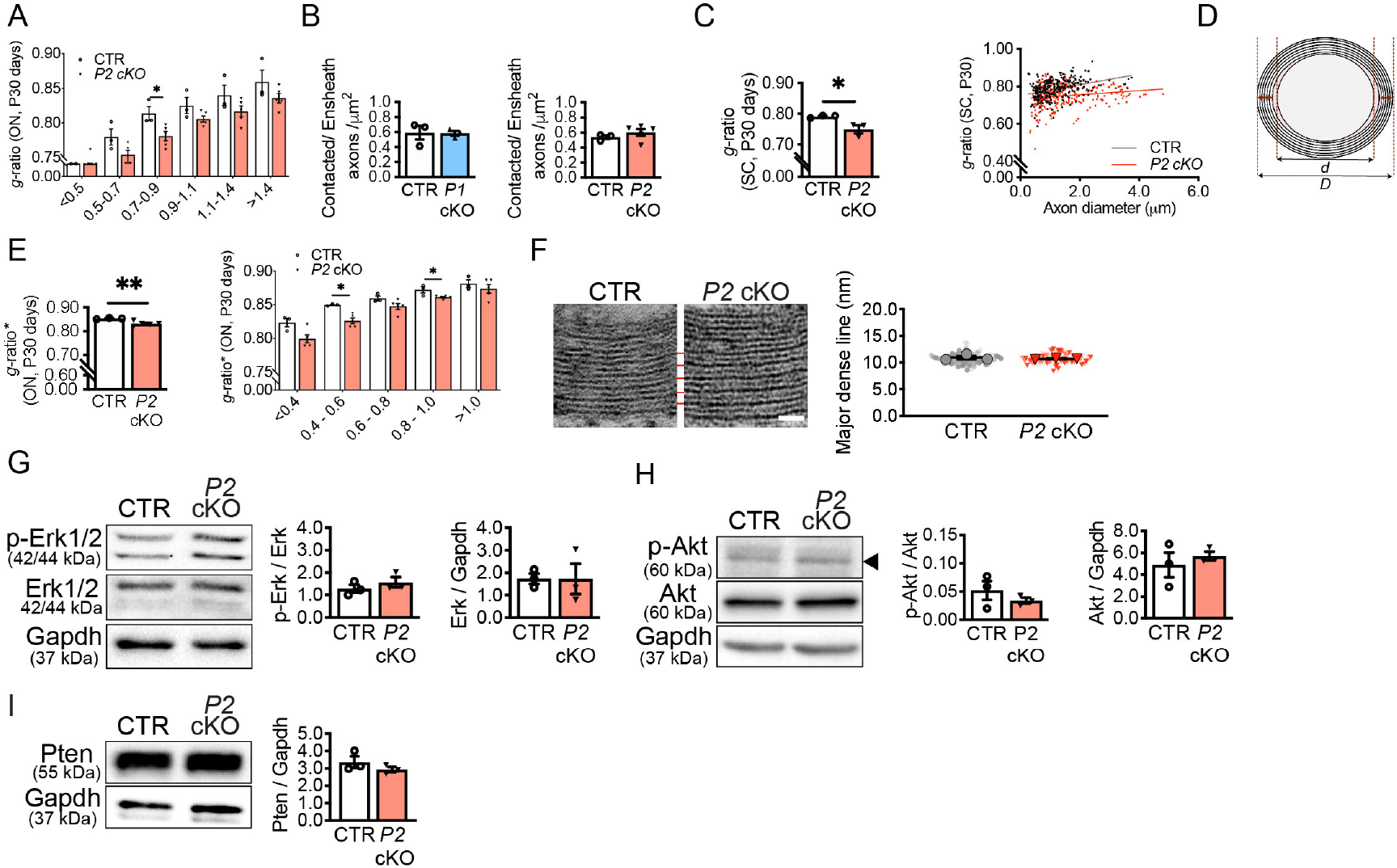
Effects on myelin structure and molecular signaling in Pinch cKOs at P30. **(A)** Distribution of g-ratios **(Fig. 3)** according to axon caliber categories. Despite the overall decrease tendency in *P2* cKO, only axons of 0.7-0.9 μm caliber are statically different. (CTR=0.813±0.010; cKO=0.781 ±0.007; *P*=0.035; two-tailed Mann-Whitney analysis of the ranks). **(B)** Similar number of myelinated axons in cross-sections of cKO and CTR at P30 (*P* = 0.36, *n*=3/5). **(C)** Lower *g*-ratios in the SC of *P2* cKO (0.750 ± 0.013 *versus* 0.781 ±0.0024, *P*=0.04). At least 92 axons were randomly quantified *per* animal (*n*=3). *G*-ratios also plotted according to axon diameter. Simple linear regression: CTR:slope=0.0285±0.0041, Y-intercept=0.753±0.0057; *P2* cKO:slope=0.01101 ±0.0038, Y-intercept=0.7343±O.o064 (*P*_slope_ =0.0029). **(D)** Schematic representation of adjusted *g*-ratio (*g*-ratio*) method (details in text). **(E)** *G*-ratio* measured in cross-section of ON from CTR and cKO **(Fig. 3).** CTR=O.854±0.OO19 *vs P2* cKO=0.832±0.004, *P*=0.009. At least 100 axons were randomly measured per animal *(n*=3/5). Distribution of *g*-ratios* according to axon caliber. Statistically significant differences found in the categories ranging from 0.4-0.6 μm (CTR=0.850±0.0009; cKO=0.826±0.001; *P*=0.036) and from 0.8-1.0 μm in diameter (CTR=0.872±0.005; cKO=0.861 ±0.001; *P*=0.04). Differences within ranks analyzed by Mann-Whitney non-parametric test. **(F)** Similar distance between consecutive major dense lines (red line) in myelin from CTR and *P2* cKO (P90). Scale bar: 20 nm. (P= 0.53, *n*=3 (at least 75 axons analyzed *per* genotype)). **(G-I)** Western blot analyses show no variations in phospo-Erk1/2 (*P*= 0.76), Akt (Thr308) (*P*=0.36) and Pten (*P*= 0.37) in ON. Statistical analysis used unpaired two-tailed Student’s *t*-test **(B, C, F, G-l).** Bars represent means±s.e.m. **(A-C, E, G-I)**.

**Fig. S5.**
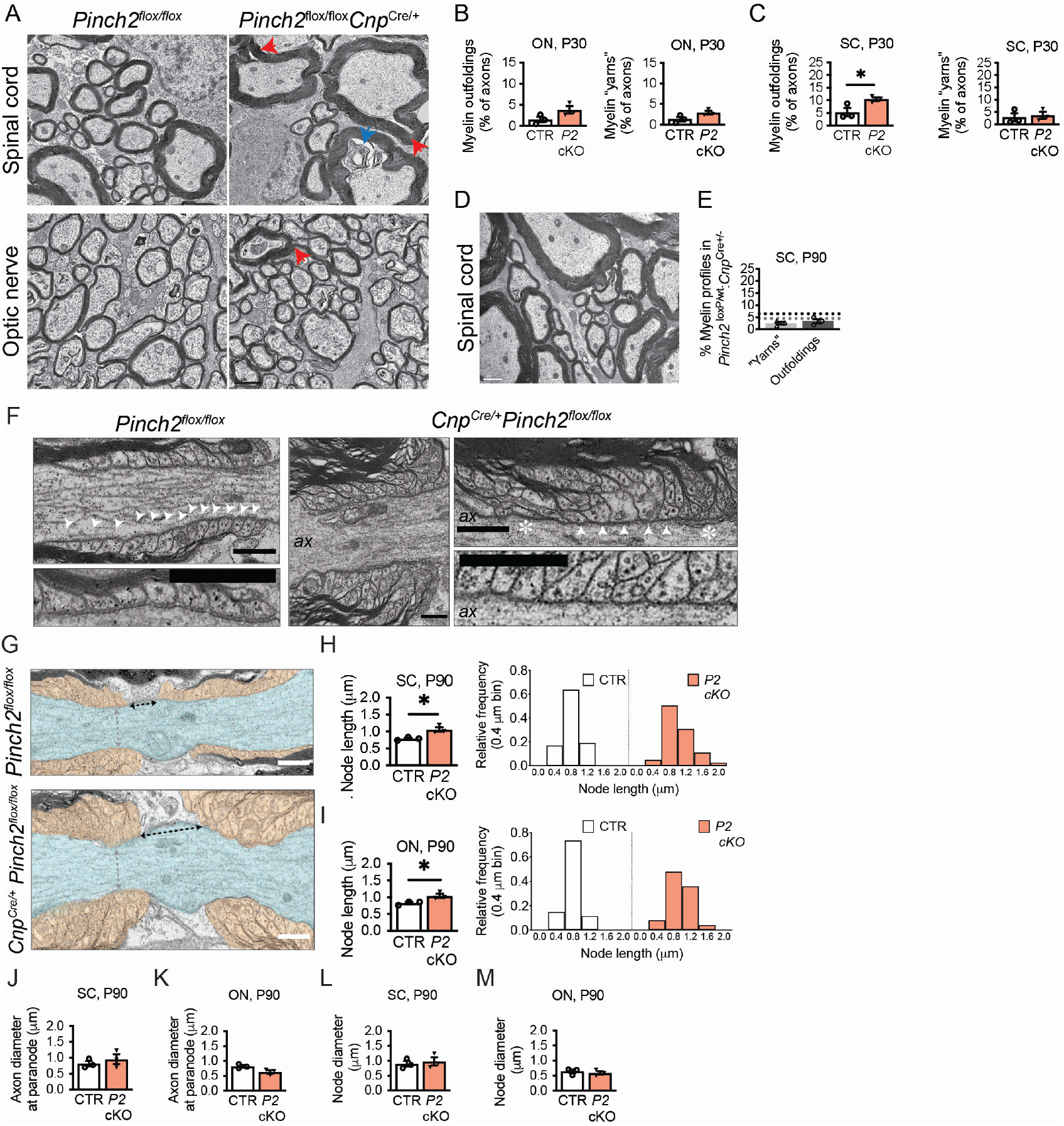
Altered myelin ultrastructure and disruption of node architecture in *Pinch2 cKO*. **(A)**Representative electron micrographs of cross-sections of optic nerve (ON) and spinal cord (SC) from conditional *Pinch2* KO (*Pinch2*^flox/flox^ *Cnp*^*Crel+*^, *P2* cKO) and control (*Pinch2*^flox/flox^, CTR) mice at P30 (red arrowheads are myelin outfoldings; blue arrows: other myelin redundant structures (’’whorls”)). Scale bars: 1 μm. **(B,C)** Despite the overall increasing tendency, the occurrence of myelin outfoldings is significant only in SC at P30 (5.37±1.67 *versus* 10.50±0.70, *P*=0.047, *n*=3). **(D)** Representative electron micrograph of a cross section of a SC from *Cnp*^*Cre,+*^ *Pinch*2f^flox/wt^. **(E)** Quantification of myelin profiles, using same criteria as in **(C)** and in **Fig. 3C**, show that whorls and outfoldings numbers are independent of *Cre* expression (*n*=3). **(F)** Longitudinal images showing alterations at the axoglial junctions at paranodal regions (SC, P90). Scale bars: 0.5 um. ax: axon; arrow heads:loops; asterisk: changed structure of paranodal elements. **(G)** Longitudinal EM sections of spinal cord (SC) of a 90 days-old *P2* cKO mice and a control littermate. Scale bars: 200 um. (H,I) Node length is increased in SC (*P*=0.018, *n*=3) and ON (*P*=0.033, *n*=3) from *P2* cKO at P90. Histograms show the differences in the distribution of node lengths in SC and ON. **(J,M)** Alterations in the nodes of Ranvier do not impact on axon diameter (*P*=0.45 in SC and *P*=0.05, *n*=3,) nor on node diameter (*P*=0.63 in SC and *P*=0.59, *n=3* in ON). Myelinated axons analyzed from **(H,I).** Statistical analysis used unpaired two-tailed Student’s Mest-Bars represent means±s.e.m. **(B,C, E, H-M)**.

**Fig. S6.**
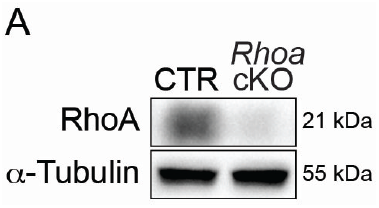
*Cnp*-driven recombination of *floxed Rhoa*alleles. **(A)** Western blot analysis of RhoA in *Rhoa* cKO (*Cnp*^*Cre/+*^ *Rhoa*^*flcx/flcx*^, *Rhoa* cKO) and control (*Rhoa*^*flcx/flcx*^, CTR) mice at P25.

**Fig. S7.**
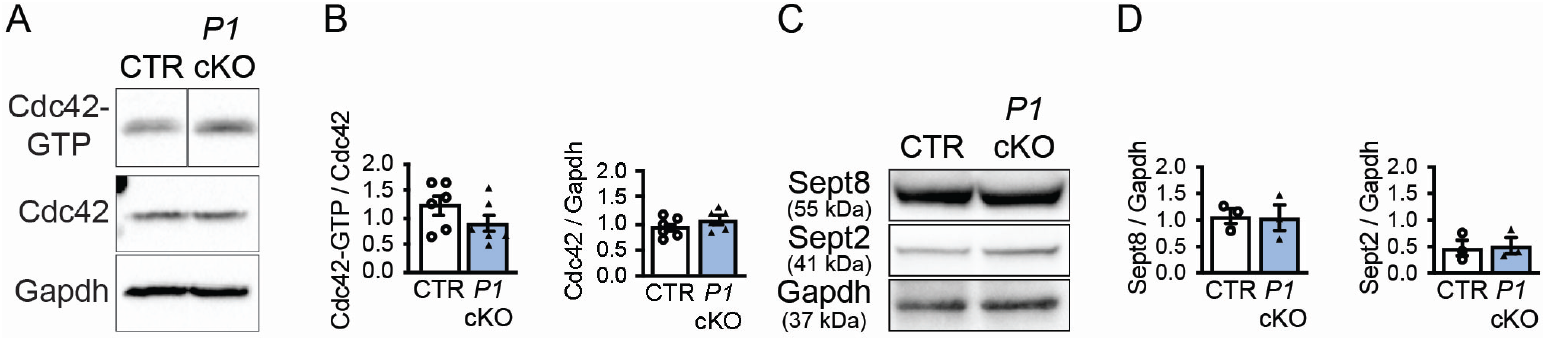
Cdc42/Sept axis is unchanged in *Pinchl* cKO. **(A)** Pull-down assay for Cdc42 in ON lysates of *P1* cKO and CTR. (**B**) Quantification of the activation index (normalized by input) shows no alterations in the active form of Cdc42 (*P* = 0.20, *n*=6) or in the expression levels (*P*=0.23, *n*=6). Each *n* corresponds to 4 pooled nerves. **(C,D)** No variations observed in the expression of Septins 8 and 2 in *P1* cKO (*P*=0.93, Sept8 and *P*=0.90, Sept2). Statistical analysis used unpaired two-tailed Student’s Mest (*n*=3) **(B, D)**. Bars represent means±s.e.m. **(A, C)**.

